# The Accuracy of Histopathological and Cytopathological Techniques in the Identification of the Mycetoma Causative Agents

**DOI:** 10.1101/489286

**Authors:** Emmanuel Edwar Siddig, Najwa Adam Mhmoud, Sahar Mubarak Bakhiet, Omnia Babekir Abdallah, Salwa Osman Mekki, Nadia I El Dawi, Wendy Van de Sande, Ahmed Hassan Fahal

## Abstract

Mycetoma is a devastating neglected tropical disease, caused by various fungal and bacterial pathogens. Correct diagnosis to the species level is mandatory for proper treatment. In endemic areas, various diagnostic tests and techniques are in use to achieve that, and that includes grain culture, surgical biopsy histopathological examination, fine needle aspiration cytological (FNAC) examination and in certain centres molecular diagnosis such as PCR.

In this retrospective study, the sensitivity, specificity and diagnostic accuracy of grain culture, surgical biopsy histopathological examination and FNAC to identify the mycetoma causative organisms were determined. The histopathological examination appeared to have better sensitivity and specificity.

The histological examination results were correct in 714 (97.5%) out of 750 patients infected with *Madurella mycetomatis*, in 133 (93.6%) out of 142 patients infected with *Streptomyces somaliensis*, in 53 (74.6%) out of 71 patients infected with *Actinomadura madurae* and in 12 (75%) out of 16 patients infected with *Actinomadura pelletierii*.

FNAC results were correct in 604 (80.5%) out of 750 patients with *Madurella mycetomatis* eumycetoma, in 50 (37.5%) out of 133 *Streptomyces somaliensis* patients, 43 (60.5%) out of 71 *Actinomadura madurae* patients and 11 (68.7%) out of 16 *Actinomadura pelletierii*. The mean time required to obtain the FNAC result was one day, and for the histopathological examinations results it was 3.5 days, and for grain, it was a mean of 16 days.

In conclusion, histopathological examination and FNAC are more practical techniques for rapid species identification than grain culture in many endemic regions.

**Author summary:** In mycetoma endemic regions, the medical and health settings are commonly suboptimal, and only a few diagnostic tests and techniques are available. That had badly affected the patients’ proper diagnosis and management and thus the late presentation of patients with advanced disease. In this retrospective study, the experience of the MRC on the common in use diagnostic tests in the period between 1991 and 2018 is presented.

In this study, the sensitivity, specificity rates and diagnostic accuracy of grain culture, surgical biopsy histopathological examination and FNAC to identify the mycetoma causative organisms were determined. The histopathological examination appeared to have better sensitivity and specificity. Furthermore, the grain culture identification needs high experience, it is the tedious procedure, and cross-contamination is common hence misdiagnosis is frequent. It can be concluded that histopathological examination and FNAC are more practical techniques for rapid species identification than grain culture in many endemic regions with poor diagnostic setting.

## Introduction

Mycetoma is a chronic granulomatous subcutaneous inflammatory infection endemic in subtropical and tropical regions, but it is reported globally [1,2]. It is characterised by a painless subcutaneous swelling, multiple sinuses formation and a discharge that contain grains [3,4]. The clinical presentation can give a clue to the diagnosis, but without further diagnostic testing it will to misdiagnosis and inaccurate treatment [5]. Mycetoma can be caused by different bacteria (actinomycetoma) or fungi (eumycetoma) [6.7]. More than 70 different micro-organisms were reported to cause this infection, and hence it is essential to identify the causative agents to the species level for appropriate treatment [8,9]. In endemic regions, the most commonly used tools are culturing of the grains, surgical biopsy followed by histopathological examination and fine needle aspiration cytological (FNAC) examination [10,11].

Currently, culturing the grains culture is still considered to be the golden standard for species identification in many centres [12,13]. However, this technique is tedious, time-consuming due to the slow growth rate and it needs expert microbiologists to identify the causative agents based on the macroscopic appearance of the isolates. Furthermore contamination is common. Patients on medical treatment may have non-viable gains, and hence it is difficult to identify the causative organism [14,15].

To overcome these difficulties, histological examination is often used complementary to culture. In a histopathological examination, it is easy to discriminate between fungal and bacterial causative agents [16,17]. However, identification to the species level is more challenging and considered far from reliable [18.19]. At the Mycetoma Research Centre (MRC), University of Khartoum, Khartoum, Sudan FNAC is a common tool to identify the causative organisms. It is less invasive, and time-consuming compared to the histopathological and culture techniques [20,21]. However, to the best of our knowledge there was no study performed in which the sensitivity and specificity of the three techniques for the identification of the mycetoma causative organisms were compared. With this background, this study was conducted at the Mycetoma Research Centre were 8500 confirmed mycetoma patients were seen and treated. In this retrospective study, the records of these patients were reviewed, and patients who undergone the three diagnostic tests were included.

## Materials and Methods

### Study cohort

Following the Mycetoma Research Centre Institutional Review Board ethical approval, all the histopathological, cytological and microbiological reports of the patients seen in the Mycetoma Research Centre over a 27-year period (January 1991 to January 2018) were reviewed. The data were collected in the pre-designed data collection sheet. The data analysis was anonymised The patient demographic characteristics, results of the three techniques were collected.

In this study, only patients in whom the causative organisms were identified by culture and had undergone both a fine needle aspirate for cytological examination and deep-seated excisional biopsy for histopathological examination were included. (Fig. 1).

**Fig 1:**
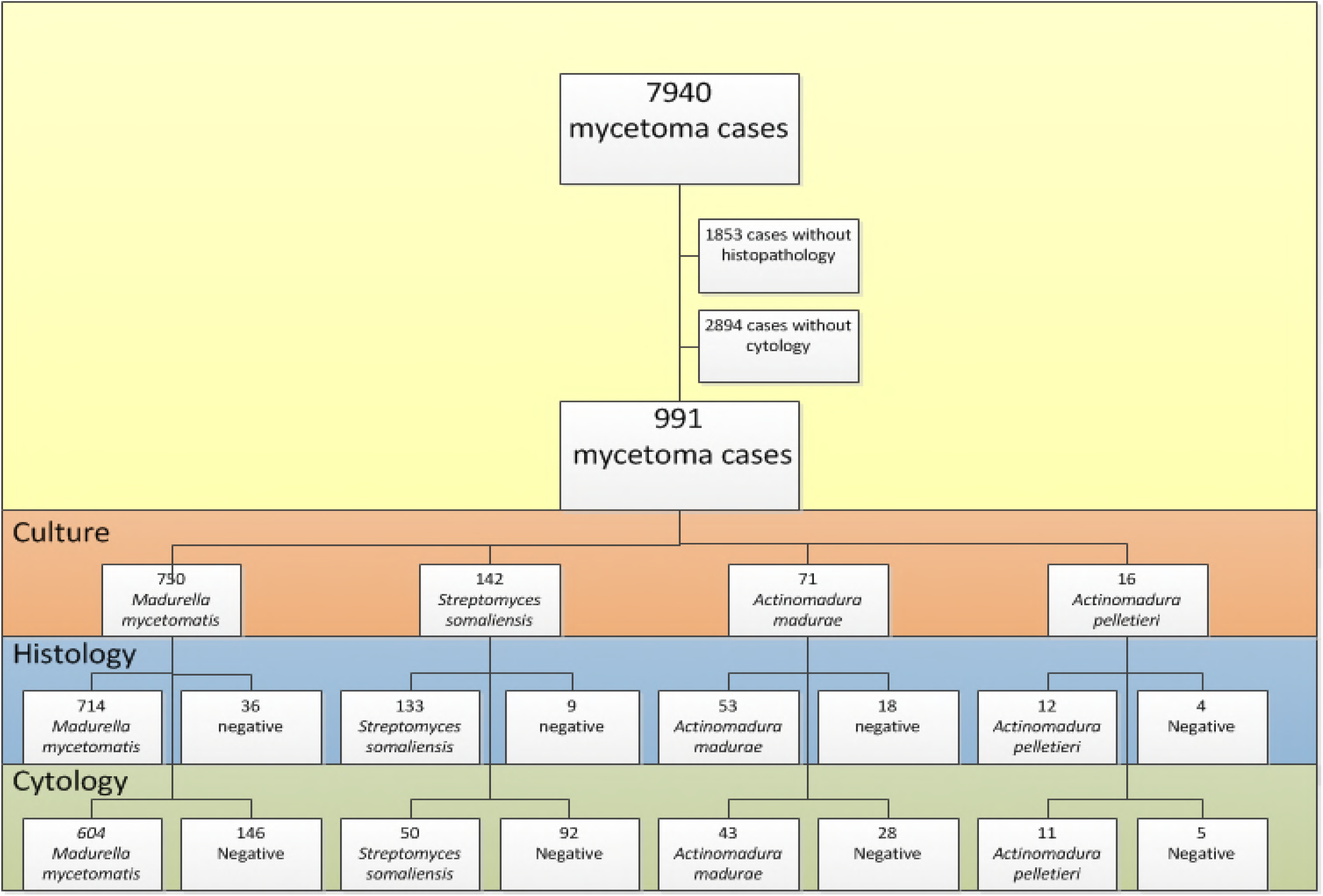
the study data collection method

The diagnostic accuracy of these techniques for identification of the causative agents was calculated as the percentage of cases with culture identification divided by the number of cases with species identified by histopathological or cytological techniques.

### Grains isolation

The grains were obtained by surgical biopsy and/or FNAC. For the latter, a 25-gauge needle was inserted into the lesion, and aspirates were taken. The yield of grains was assessed visually by the number and size of grains obtained. If the yield was low, a second aspiration was taken with a 23-gauge needle. When excessive bleeding from the lesions was encountered, a 27-gauge needle was used. The obtained sample usual spiltted into two parts; one was transported immediately to the microbiology department for culturing, the other part was sent to the histology department for histopathological and cytological.

### Grains culture

The mycetoma grains were washed three times in sterile normal saline and cultured on Sabaroud dextrose agar with gentamicin for fungal grains. The actinomycetoma gains were cultured in Yeast Extract agar. Grains were incubated at 37°C and growth was checked daily. The isolates were identified by their microscopical appearance and biochemical testing.

### Cytological Examination

The aspirate was allowed to air dry and was stained using Diff-Quick stain. The stained aspirates were examined by an expert histopathologist for the presence of the following cytomorphological features: smears cellularity, the host inflammatory tissue reaction, the presence and types of the causative organisms’ grains. Species identification was based on species-specific criteria. In general, *M. mycetomatis* grains can be either small or large, are light to dark brown in colour and have irregular outlines and a crushing artefact when stained with hematoxylin and eosin (H&E) (Fig. 2A).

**Fig 2:**
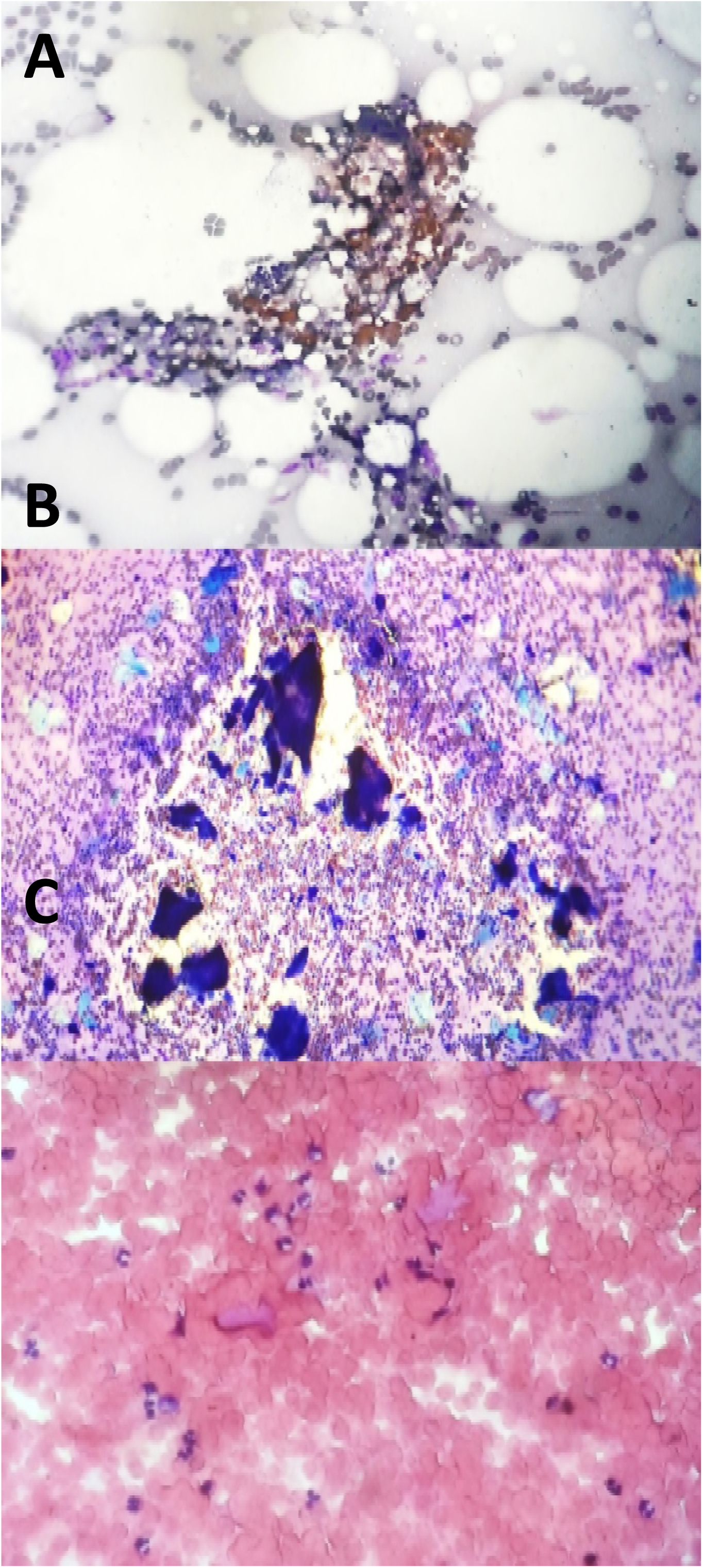
photomicroscopy of the cytopathological appearance of some mycetoma causative agents. *M. mycetomatis* (A), *A. pelletierii* (B), *S*. somaliensis in fine (H&E X10)

S. *somaliensis* grains are difficult to see in H&E stained sections, they stain bright pink to hazy pink in colour, are often oval to irregular shaped and can be as aggregates (Fig. 2C).

*A. madurae* grains are small oval shaped, and it stained pink to red colour in H&E and tend to be as one mass without any fractures. *A. pelletierii* grains are small rounded to oval shaped, and they stained deep blue in H&E stained sections and tend to be fractured.

### Histopathological Examination

All patients underwent surgical biopsy under anaesthesia, which was fixed in 10% formalin and processed further into paraffin blocks. 3-5-μm sections were obtained and stained with H&E. Species identification was made based on species-specific criteria. *M. mycetomatis* grains tend to be large, light to dark brown in colour with irregular outlines. They tend to fracture when sections are cut. *M. mycetomatis* has two different types of grains, and these are the filamentous and vesicular. The filamentous type, is the most common type and consists of brown septated and branched hyphae that may be slightly more swollen towards the edges (Fig. 3D).

**Fig 3.**
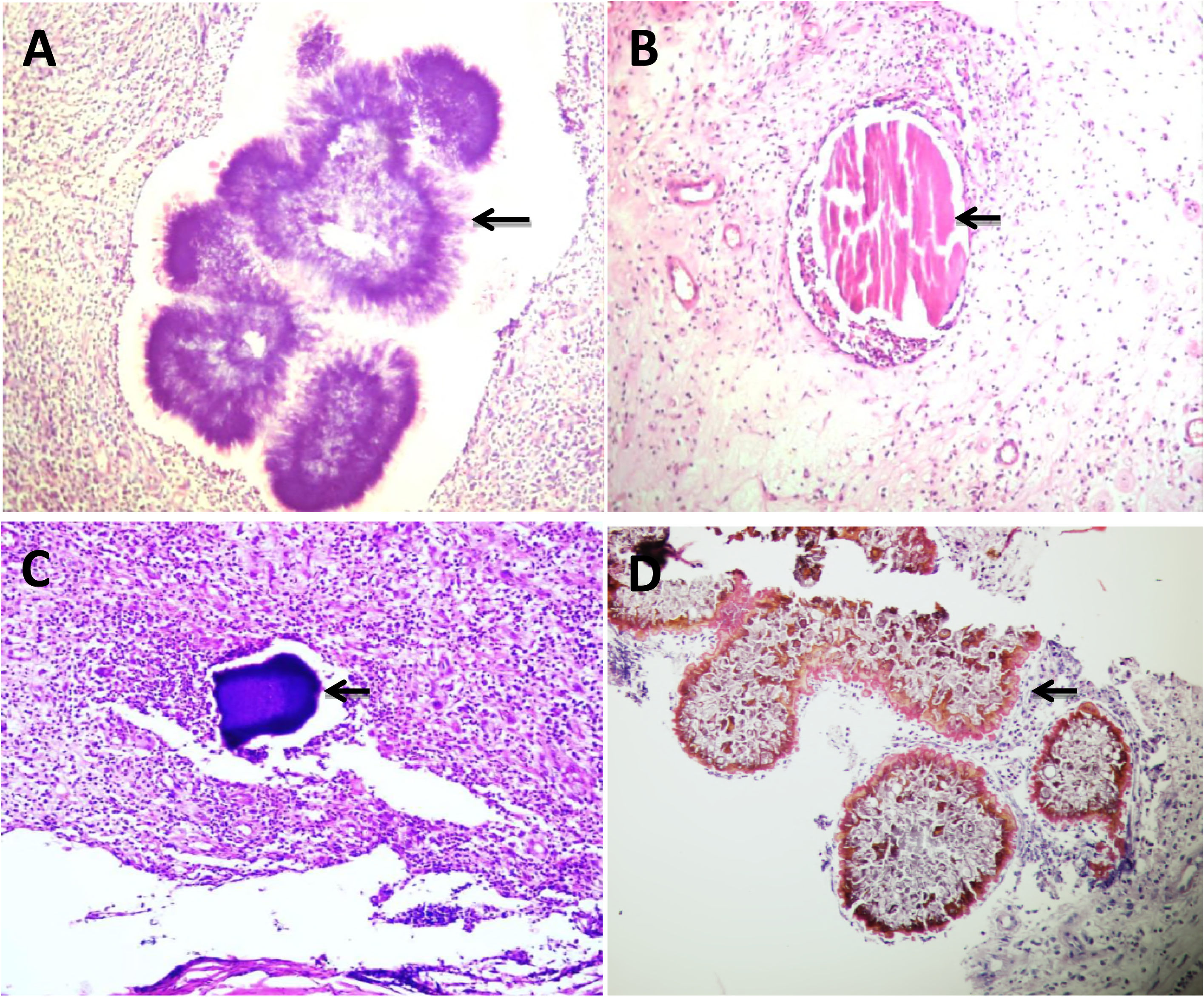
Histopathological appearance of mycetoma causative agents; *A. madurae* (A), *S. somaliensis* (B), *A. pelletierii* (C) and *M. mycetomatis* (D). (H&E X10).

*S. somaliensis* grains are rounded to oval in shape, with homogenous appearance in tissue sections. They appear faint yellow in unstained sections, and the grains are not well stained with H&E. Moreover, as a result of sectioning they may show longitudinal cracks, the filaments are fine (measured between 0.5 – 2 μm in diameter), closely packaged and embedded in cement matrix (Fig. 3B).

*A. pelletierii* grains are small, round to oval in shape and semicircular and sickle like shapes have been observed as well. The filamentous structures are pretty difficult to be detected. However, a careful and meticulous examination of the periphery of the grains may show some of them. *A. pelletierii* grains stain deep violet with H&E, which is very characteristic and allows the definitive diagnosis without a need for culturing techniques (Fig. 3C).

*A. madurae* grains ranged from yellow to white. Therefore, it can be difficult to discriminate them from the surrounding fat. Histologically the grain size ranges from small to large. The large grains have a characteristic variegated pattern. The periphery of the grain is opaque, homogenous and deep purple when stained with H&E stain, while the centre is less densely stained. Additionally, the periphery of the grains shows an eosinophilic material (Fig. 3A). Smaller grains are more homogeneous and are difficult to distinguish from *A. pelletierii*. However, even the small grains of *A. madurae* have a more deeply stained purple fringe, which is not seen in *A. pelletierii*.

## Results

In this study, 991 patients out of 7940 patients were eligible and were included in the analysis. Their ages ranged between 5 and 75 years old. The majority were males 737 (74.3%), and most of them were students 327 (32.9%) and farmers 167 (16.8%). The majority of the patients (837 out of 991), gave a history of discharge that contained grains and the majority of these grains were black (565; 57%)) followed by yellow (104;10.5%), white (60; 6.1%) and red grains (14; 1.4%). In this cohort, the majority of patients, (72.6%) had no history of local trauma, only 191 (19.3%) patients did recall a local trauma and the remaining 73 (7.4%) patients were not certain.

Based on the culture reports of the grains, in 750/991 (75.6%) of the patients the mycetoma was caused byM. *mycetomatis*, in 142/991 (14.4%) it was caused by S. *somaliensis*, in 71/991 (7.16%) it was caused by *A. madurae* and in 16/991 (1.6%) it was caused by *A. pelletieri*. In 11 patients no growth was reported from the grains obtained during the sample collection. The time to growth differed case by case and ranged between 5 and 28 days.

In this study, out of the 991 mycetoma cases, the correct species identification was obtained for 912 cases using histopathological examination. Using FNAC, the correct diagnosis was obtained in 708 cases. The histopathological examination confirmed the diagnosis of *M. mycetomatis* in 714 of 750 cases with 95.2% sensitivity, 95.4% specificity and diagnostic accuracy of 95.3%. For FNAC only 604 out of 750 *M. mycetomatis* cases were identified, resulting in a sensitivity of 80.5%, a specificity of 88.4% and a diagnostic accuracy of 82.4%.

Out of 142 *S. somaliensis* cases, 133 were also identified with histopathological examination with 93.7% sensitivity, 98.9% specificity and diagnostic accuracy of 98.2%. With FNAC only 50 out of 133 S. *somaliensis* cases were identified, resulting in a sensitivity of 35.2%, a specificity of 99.3% and a diagnostic accuracy of 90.1%. 53 out of 71 cases with *A. madurae* identification were identifuied by histopathological examination, with a sensitivity of 74.7%, 99.5 % specificity and diagnostic accuracy of 97.7%. FNAC identified 43 out of 71 cases with a sensitivity of 60.6%, specificity of 94.4 % and diagnostic accuracy of 91.9%.

For *A. pelletierii* out of 16 cases; 12 were also identified with histological examination with 75.0% sensitivity, 100% specificity, and diagnostic accuracy of 99.6%. For FNAC a sensitivity of 68.8%, a specificity of 99.7% and a diagnostic accuracy of 99.2% were obtained.

With the histopathological examination, false negative result was reported in 36/750 *M. mycetomatis* cases, 9/142 *S. somaliensis* cases, 18/71 *A. madurae* cases and 4/16 *A. pelletierii* cases. To determine the false negative results reasons, the histopathological slides were re-examined. There were various reasons for the false negative, and that included the absence of mycetoma histopathological architecture resulted in overlooking the causative agent (Fig 4). Furthermore, in some blocks, the grains were absent, either because the tissue was not homogenously infected by the causative agent and that the part which was taken for histology or the section contained no grains.

**Fig 4:**
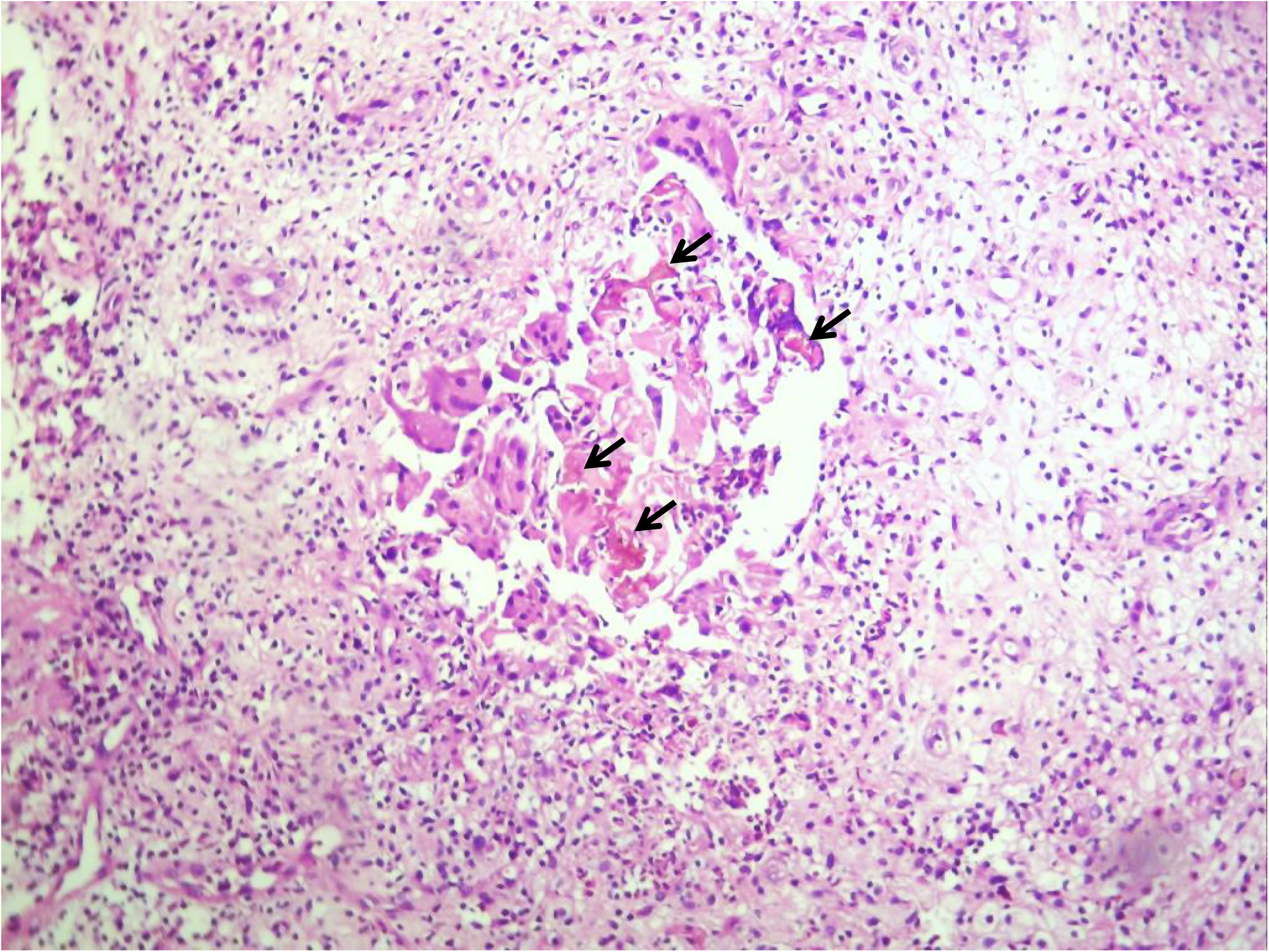
photomicroscopy showing histopathological section with histiocytes at the centre with sparse amount of *M. mycetomatis* the grains (arrows). It was reported first as false negative. (H&E X 10).

This latter might be overcome by examining multiple sections at different depths of the histology blocks especially when inflammation and necrosis are noted.

False positive results were obtained in 28 of the cases. This was attributed to the presences of numerous structures that can mimic the appearance of *M. mycetomatis* and that included vegetables, synthetic fibres and algae (Fig 5) which can resemble fungal hyphae and calcification (Fig 5.). In overall, using histology correct species identification was obtained in the majority of cases.

**Fig 5:**
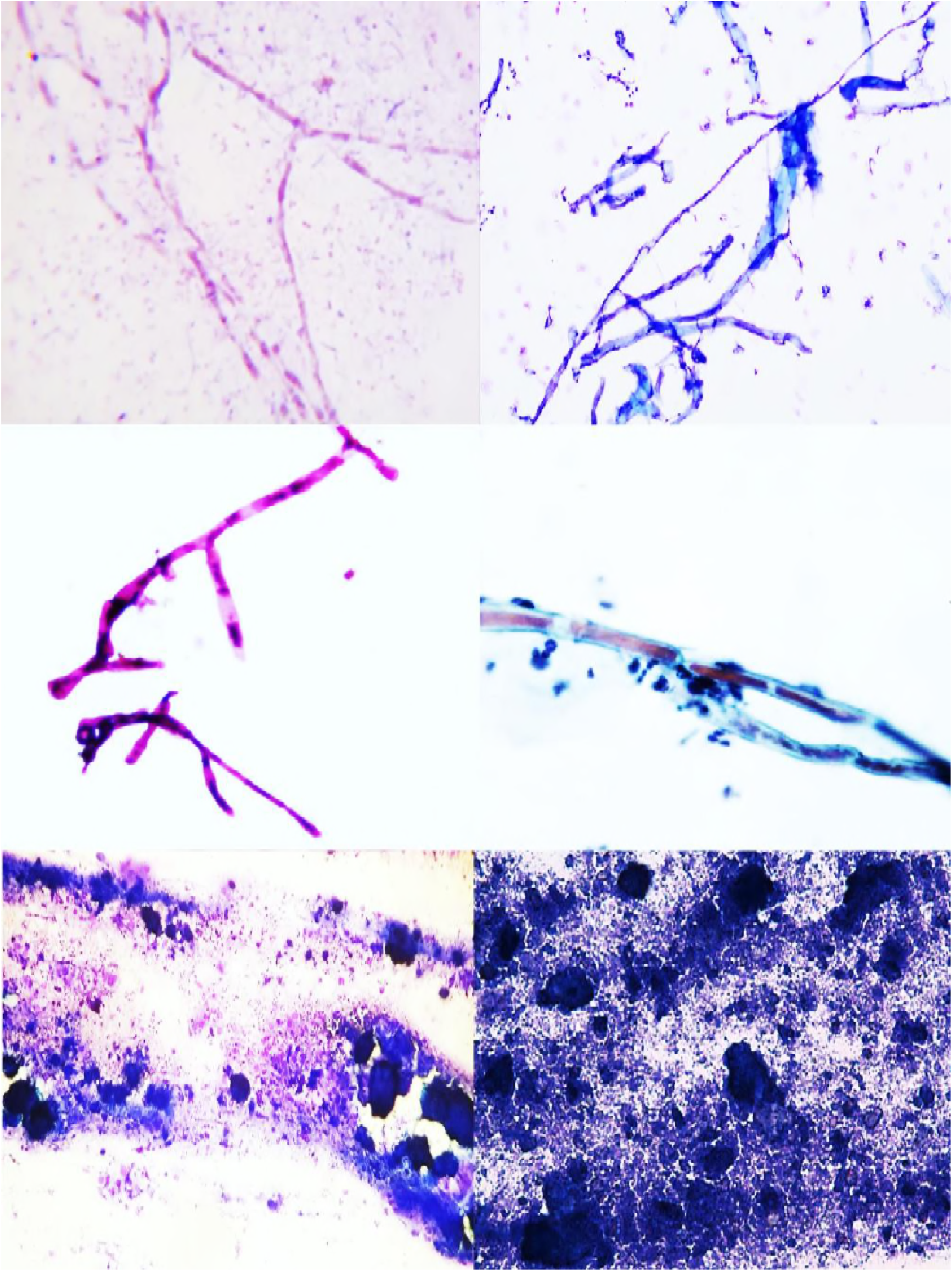
false positive identifications in cytology. Showing cytological smears of (A) *M. mycetomatis* hyphae stained with Giemsa stain, (B) Synthetic Fibers stained Wright-Giemsa stain. (Giemsa stain, X40). (C) Smear showing hyphae of *M. mycetomatis*. (D) Smear showing elongated structures of the Oedogoniales order. The chloroplasts form a chain interrupted by clear zones (X40). (E) Smear showing *M. mycetomatis* grains after being crushed on the smear. (F) smear with abundant calcific debris without intact cells taken from patient with tumoral calcinosis (Diff Quick, X10).

The mean time to identify the culture isolates was 16 days (range 5 to 28 days), for histology it was 3.5 days (range 2 to 5 days), and for cytology, it was one day (range 1 to 2 days). This demonstrated that reliable species identification using histology was obtained in 92.0% of cases within an average time reduction of 13.5 days, for cytology this was 71.4% of cases with time reduction of 15 days, indicating that adding histology or cytology to the diagnostic techniques used for species identification resulted in an earlier start of treatment.

## Discussion

The accurate identification of mycetoma causative agents is considered the cornerstone for the initiation of appropriate therapy. Hence a rapid and accurate diagnostic tool to achieve the definitive species identification is considered a critical part in patient treatment and management [21–23]. Different laboratory techniques for species identification are in use, including culture, histophatology, [7, 25], FNAC [8, 24], serological assays and imaging [26 – 29] as well as different molecular diagnostic tools [30 – 35]. However, not all these assays are available in endemic regions. In the Mycetoma Research Centre, culturing of the grains, histopathology and FNAC are routinely performed and have been used for the past 27 years. In this communication we have used the data collected for the last 27 years to assess the sensitivity, specificity and diagnostic accuracy of histopathology and cytology in the identification of mycetoma causative agents in comparison to the current golden standard: culturing.

This study showed that the histopathology was more accurate to FNAC in terms of species identification. Our results are in line with that reported previously by Yousif and colleagues [36]. They reported 90.9% agreement when histopathology was compared to FNAC for the diagnosis of *M. mycetomatis* (90.9%) while for actinomycetoma causative agents it was only 60%. The lower diagnostic agreement of actinomycetoma causative agents could have been caused by morphological similarities of these microorganisms. Furthermore, both techniques are operator dependent and need intensive training and experience which could have its reflections on the accuracy.

Mycetoma can be caused by more than 70 different causative agents [37], but the distribution of these species is not everywhere the same which could cause differences in diagnostic accuracy in different regions. In some of the mycetoma endemic regions, mycetoma is caused by closely related species. Morphologically these organisms may look similar which could cause a challenge in the identification of these organisms. In Mexico, the most common causative agents are *Nocardia brasiliensis* and *Nocardia asteroides* [37], two closely related species which are difficult to differentiate from each other based on histopathology [38, 39]. In Senegal, the most common causative agents of eumycetoma are *M. mycetomatis* and *Falciformispora senegalensis* which both can cause black grain mycetoma [37, 40]. In the black grains of *F. senegalensis*, the centre is non-pigmented, and the cement is absent, whereas at the peripheries the grains are dark coloured and brown cement is present. However, this is also seen in black grains of *Trematosphaeria grisea* and certain grains of *M. mycetomatis*. Hence an expert pathologist is needed to differentiate between these organisms [41].

The study performed here was a retrospective study, looking back at the records of the Mycetoma Research Centre for the past 27 years. During that time molecular identification of the causative agents was not performed and culture was considered the golden standard. Recently in the study conducted by Borman and colleagues demonstrated that using morphological identification, misidentifications occurred in many cases [42]. Out of 28 previously identified *Trematosphaeria grisea* isolates, 22 were, other fungal species [42]. For actinomycetoma causative organisms, misidentifications also have been described. In 2008, Quintana and associates demonstrated that half of the S. *somaliensis* isolates obtained from Sudan appeared to be *Streptomyces sudanensis* [43]. Furthermore, next to *A. madurae* and *A. pelletieri* also *Actinomadura latina* was described [44].

In this study, the sensitivity of histopathological technique was superior to that FNAC for all species tested. In that study they studied the performance of FNAC in comparison to histology in 19 different mycetoma patients. Out of these 19 patients, five patients had to be excluded due to inadequate aspirated materials. From the 14 remaining patients, 10 were diagnosed as *M. mycetomatis* with histopathology, and 4 were actinomycetoma. With this limited number of patients, they could conclude that FNAC could identify the causative agent in 9 out of 10 *M. mycetomatis* patients. One patient identified by histology could not be identified with cytology, again confirming that histology was superior to FNAC in respect to species identification [15]. A result confirmed in our current study, as in our study 146 patients with *M. mycetomatis* mycetoma were missed with FNAC. However, of the three different identification methods used, FNAC was the most rapid and resulted in a species identification within 1 day, instead of 3.5 days for histology or 16 days for culture. FNAC is a simple and rapid diagnostic technique which can be used at the one-stop diagnosis clinic and in epidemiological and field surveys. However, it has many limitations: it is an operator dependent technique, can be painful and can lead to deep-seated bacterial infection. At the moment the fine needle aspirate is often taken blindly without guidance of ultrasound imaging which creates a risk that the operator might miss the pockets which contains grains. With the use of the ultrasound-guided aspiration, the diagnostic yield of the technique will improve which in its turn could enhance the number of cases in which a positive species identification might be obtained.

The grains culture remains in many centres the cornerstone for the diagnosis of mycetoma. However it is a time-consuming procedure and an experienced microbiologist is needed to identify the organisms to the species level. Cross-contamination is a common problem. However, by complementing culturing with histology or FNAC a preliminary identification might be obtained earlier.

